## Supporting Information for "Single molecule delivery into living cells"

### Methods

All chemicals, unless otherwise specified, were obtained from Fisher Scientific at ACS grade.

#### Cell culture

All cells were cultured in cell culture flasks (83.3911.302; Sarstedt) or 35 mm glass bottom dishes (FD35-100; World Precision Instrument Ltd) inside an incubator at 37°C with 5% $CO_2$.

Table 1 lists the cells used in this study and the culture medium used.

**Table 1. Cell lines used in this study and the culture medium composition.**

| Cell Line | Origin | Culture medium |
|---|---|---|
| HeLa | CVCL_0030, Verified by ECACC | Dulbecco's Modified Eagle's Medium (D5671; Sigma Aldrich) supplemented with 1X GlutaMax™ (35050038; Thermo Fisher), 1X Penicillin Streptomycin (15140122; Fisher Scientific) and 10% (v/v) foetal bovine serum (F7524; Sigma Aldrich) |
| HeLa RNuc | Established by expressing pmCherry-NLS in the original HeLa CVCL_0030. Followed by FACS sorting | Dulbecco's Modified Eagle's Medium (D5671; Sigma Aldrich) supplemented with 1X GlutaMax™ (35050038; Thermo Fisher), 1X Penicillin Streptomycin (15140122; Fisher Scientific) and 10% (v/v) foetal bovine serum (F7524; Sigma Aldrich) |
| Human Umbilical Vein Endothelial Cell (HUVEC) | C-12253; Promocell | Endothelial Cell Growth Medium (C-22010; PromoCell), 1X Penicillin Streptomycin (15140122; Fisher Scientific) |
| Primary Rat Cortical Neurons | A36511; Gibco | Neurobasal medium (21103049; Gibco), 1X GlutaMax™ (35050038; Thermo Fisher), 1X B-27 supplement (17504044), 1X Penicillin Streptomycin (15140122; Fisher Scientific) |
| Primary Mouse Dorsal Root Ganglion (DRG) Neurons | Provided by Prof Nikita Gamper's research group in University of Leeds | Neurobasal medium (21103049; Gibco), 1X GlutaMax™ (35050038; Thermo Fisher), 1X B-27 supplement (17504044), 1X Penicillin Streptomycin (15140122; Fisher Scientific) |

For dividing cells (HeLa, HeLa RNuc, NHEK and HUVEC), the cells were passaged when they were confluent. The cells were washed with 1X PBS (D8537; Sigma Aldrich), followed by the addition of a pre-warmed 1X trypsin-EDTA (T3924; Sigma Aldrich), the cells were then left inside the incubator for 3 minutes, the detached cells were collected by centrifugation at 500Xrcf to remove the trypsin-EDTA, the pellet was immediately resuspended in culture medium, then plated into a cell culture flask or onto the 35 mm glass bottom dish. To cryopreserve the cells for long term storage, after the removal of the trypsin-EDTA solution, cells were resuspended in Synth-a-Freeze™ Cyropreservation medium

(A1254201; Gibco) at minimum of 1X10$^6$ cells/ml. A minimum of 100 µl of the resuspended cell mixture was dispensed into a 1.8 ml cryovial (E3090-6222; Starlab). The cryovial was immediately sealed inside a polystyrene box and frozen down overnight at -80°C, and then transferred to a liquid nitrogen canister for long term storage.

For neuronal cell-culture, the glass bottom dish was pre-coated with synthetic laminin peptide (SCR127; Sigma Alrich) according to the manufacturer's instructions. After treatment, all solutions were removed from the inside of the dish and rinsed with $ddH_2O$, then the dishes were dried inside the flow hood and wrapped with parafilm and stored at 4°C. Prior to use, the dishes were rinsed two times with pre-warmed culture medium. For the primary rat cortical neurons, the 1 ml thawed cell vial (1X10$^6$ cells) was mixed with 11 ml of the pre-warmed culture medium and plated equally onto six laminin coated dishes. For the primary mouse DRG neurons, the cells were obtained by dissection carried out by Dr Vincenzo Prato of Prof Nikita Gamper's research group at the University of Leeds, the neurons were plated onto the laminin coated 35 mm glass bottom dishes. The day of either thawing or dissection was designated as day 0 and all neurons were nanoinjected between 3 to 10 days.

#### Cell transfection and establishment of the HeLa RNuc stable cell line

Three plasmids were used to transfect cells in this study: pMaxGFP, pmCherry-NLS and the pSV-β-Galactosidase Control Vector (E1081; Promega). To transfect cells, Lipofectamine® 2000 (11668030; Thermo Fisher) was used. The cells were plated at 50% confluency in a 35 mm dish prior to the day of transfection in standard culture medium. 4 µg of the plasmid was diluted into 125 µl of serum-free culture medium in one Eppendorf tube, then 4 µl of lipofectamine reagent was diluted in 125 µl of serum-free culture medium in another Eppendorf tube, the two tubes were allowed to sit on bench for 5 minutes. Then the plasmid containing culture medium was dispensed drop by drop into the lipofectamine serum-free culture medium containing tube and incubate at room temperature for another 15 minutes.

The mixture was then dispensed onto the culture dish and incubated overnight; the culture medium was exchanged the next day. Transfection was confirmed by imaging using a Zeiss LSM880 with Airyscan (Zeiss) or with an EVOS microscope with an appropriate fluorescent wavelength light cube.

The establishment of the HeLa RNuc stable cell line involved the transfection of the pmCherry-NLS into the HeLa (CVCL_0030) cell line. Upon transfection, the cells were cultured in standard culture medium with G418 sulfate (10131035; Gibco) at 1 mg/ml concentration for 2 weeks. The cells were then sorted by fluorescence-activated cell sorting (FACS) to select for high pmCherry fluorescence. The FACS was performed by Dr Ruth Hughes at the University of Leeds Bio-imaging and Flow Cytometry facility.

#### Nanoinjection

For all nanoinjection procedures, the ion current trace was recorded by pClamp 10 (Molecular Devices). All fluorescent images were captured by the ANDOR iQ3 live cell imaging system with appropriate excitation laser and emission filter combinations. The nanopipette was lowered down at 10 μm/s during cell penetration.

Different molecules were injected and their solution prepared as outlined as below:

For fluorescein-conjugated dextran 70,000 g/mol (11520226; Fisher Scientific), the dextran was dissolved and diluted in 0.22 μm filtered 1X PBS, aliquoted and then stored at the concentration of 140 μM at -20°C. For nanoinjection, the dextran was further diluted to 140 nM in 1X PBS and the resultant solution of dextran was then used to fill =the nanopipette.

For pMaxGFP, the cells were plated onto a 35 mm glass bottom gridded dish (81148; Ibidi). The grid had lettered and numbered 4X400 squares with a 50X50 μm dimension for each square. The pMaxGFP plasmids were diluted to 1.3 nM in 1X PBS and used to fill the nanopipette. The duration of the nanoinjection part of the experiments all lasted less than 2 hours, after which the L-15 medium was replaced with the culture medium before returning to the incubator.

For β-galactosidase nanoinjection, in addition to the standard L-15 medium 2 μM SPiDER-βGal (SG02-10; DOJINDO) was added to the medium. SPiDER-βGal was the fluorescent substrate for β-galactosidase. The size exclusion chromatography purified β-galactosidase was diluted to 1 μM with 1X PBS and used to fill the nanopipette. As an injection control, a 100 nM 1X PBS diluted Alexa Fluor 594 Maleimide was also used to perform nanoinjection in the nucleus.

For the α-synuclein fibril seeds nanoinjection, the 200 μM monomeric equivalent α-synuclein fibril seeds were diluted to 1 μM monomeric equivalent with 1X PBS. The solution was then used to fill up the nanopipette for nanoinjection.

For the 7 kbp dsDNA nanoinjection and single molecule sensing experiment, the commercially available 7 kbp dsDNA (SM1741; Thermo Fisher) was diluted to 5 nM in 1X PBS with or without 10 μM ATTO 488 carboxylic acid (41051; Sigma Aldrich).

#### Measurement of β-galactosidase activity in solution

The snap frozen β-galactosidase was thawed, serial diluted and incubated with final concentration of 2.5 μM fluorescent substrate SPiDER-βGal (SG02-10; Dojindo) for 30 minutes at room temperature inside a 96-well black clear bottom microplates (3631; Corning). The fluorescent signal was measured using the CLARIOstar Plus plate reader (BMG LABTECH).

#### CTCF and fold change calculation

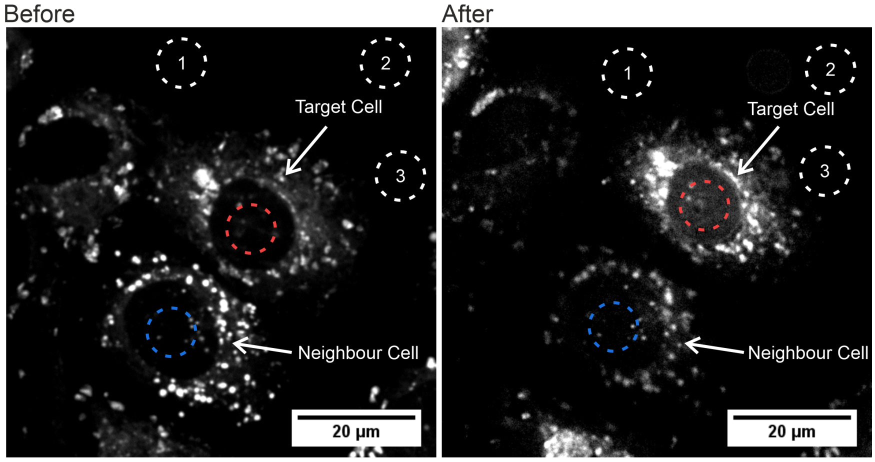


The corrected total cell fluorescence (CTCF) fold change was calculated. During the nanoinjection experiment, fluorescent images before and after nanoinjection were taken for each experiment, alongside the corresponding brightfield image of the same cell. These images were analysed with ImageJ to obtain the fluorescent signal data for the calculation of CTCF. First, three background fluorescent levels were measured by selecting three spots where there were no cells with three numbered dotted circles, which were used to calculate the average background fluorescent levels as shown in the example image above**Error! Reference source not found.**. The nanoinjected cell, denoted as the target cell here, only the fluorescent signal of the centre of the nucleus was measured, as indicated with a red dotted circle. The same measurement was performed on a cell that was not injected, denoted as neighbour cell, as shown with a blue dotted circle.

#### Generation of α-synuclein fibrils

BL21(DE3) (carries the gene for T7 RNA polymerase under control of the lacUV5 promoter) competent *E. coli* (C2527; New England BioLabs) was transformed with carbenicillin resistant pET23a encoding a codon optimised gene encoding full length human α-synuclein using the heat shock transformation. Briefly, 1 µl of 200 ng/ml plasmid was added to 50 µl of the *E. coli*, followed by incubation on ice for 30 minutes and immediately followed by heat shock procedure at 42°C for 45 seconds in a heat block. The transformed *E. coli* was then inoculated in a 25% (w/v) Luria broth medium for 60 minutes at 37°C, and then grown overnight at 37°C on 25% (w/v) Luria broth agar plates containing 25 µg/ml of carbenicillin. A single bacterial colony was picked and grown overnight at 37°C with constant shaking at 200 rpm in 150 ml 25% (w/v) Luria broth containing 25 µg/ml of carbenicillin. The bacterial culture was mixed with 30% (v/v) autoclaved glycerol at the ratio of 1:1, and aliquoted into 1 ml cryotube vials (V7884; Sigma-Aldrich) and stored at -80°C as a glycerol stock for future use.

For the large-scale expression of the α-synuclein, approximately 1 µl of the transformed bacteria glycerol stock was added to 1 ml of 25% (w/v) Luria broth containing 25 µg/ml of carbenicillin and was grown overnight at 37°C with constant shaking at 200 rpm. This starter culture was then added to 1 L of 25% (w/v) Luria broth containing 25 µg/ml carbenicillin under constant 200 rpm shaking at 37°C. Protein expression was induced by adding isopropyl β-D-1-thiogalactopyranoside (IPTG) to a final concentration of 1 mM when the culture reached an $OD_{600}$ of 0.6, followed by incubation for 4 hours.

The purification of monomeric α-synuclein was performed as following. After the IPTG induction the *E. coli* was collected by continual action centrifuge (Heraeus$^{TM}$ Biofuge$^{TM}$ Stratos$^{TM}$ Centrifuges; HCA 10.300 centrifuge rotor; Thermo Scientific) at 15,000Xrpm. The collected *E. coli* was lysed by resuspending the *E. coli* into lysis buffer (25 mM Tris-HCl, pH 8.0; 100 µg/ml lysozyme; 50 µg/ml phenylmethylsulfonyl fluoride; 20 µg/ml deoxyribonuclease (DNase); 1 mM EDTA and frozen overnight at -20°C.

The bacteria were then thawed and lysed by mechanical homogenisation followed by 10 minutes incubation at 70°C in water bath. The bacteria debris pellet was removed by centrifugation at 35,000Xrcf for 30 minutes at 4°C (JLA 16.250 rotor; Beckman). The supernatant was titrated to pH 3.5 with 1 M HCl and incubated for 30 minutes with gentle stirring using a magnetic stirrer inside 4°C cold room. Then, the supernatant was collected by centrifugation at 35,000Xrcf (JLA 16.250 rotor; Beckman) for 30 minutes at 4°C and the supernatant titrated with 1 M NaOH to pH 7.5. The solution was dialysed against 20 mM Tris-HCl pH 7.5 using a 3.5 kDa dialysis membrane (68035; Thermo Scientific) overnight at 4°C with gentle stirring constantly with a magnetic stirrer.

α-synuclein was purified from the dialysed protein solution by anion-exchange chromatography, followed by size exclusion chromatography to increase the purity. The XK 50/20 column (Amersham Bioscience) packed with Q-Sepharose (GE Healthcare) were used for anion-exchange chromatography on the AKTA Prime column purification system (GE Healthcare). The column was equilibrated with 20 mM Tris-HCl, pH 7.5 buffer (Buffer A) with 15% (v/v) of 1 M NaCl, 20 mM Tris-HCl, pH 7.5 (Buffer B). After stable $A_{280}$ and conductivity values were obtained, the protein solution was loaded onto the column and then followed by 1 column volume (approximately 300 ml) of buffer A with 15% (v/v) of buffer B, this is equivalent to a total of 150 mM of NaCl throughout the entire column. Proteins were eluted under constant gradient of 3 column volume of buffer B from 15 to 50%, i.e. 150 mM to 500 mM NaCl. Eluted fractions were collected at between 30 to 37% of the buffer B, i.e. 300 to 370 mM NaCl concentration. The collected anion-exchange fractions were then further purified by size exclusion chromatography. The eluted fractions were pooled and stored at 4°C for short term storage. The Superdex 75 10/300 GL column (GE Healthcare) was used with the AKTA Prime column purification system (GE Healthcare) and equilibrated with buffer A. The protein solution was loaded onto the column and the α-synuclein containing fractions were collected and pooled together, the fractions were dialysed against 30 mM ammonium bicarbonate using a 3.5 kDa dialysis membrane at 4°C with gentle stirring using the magnetic stirrer overnight. The protein was then lyophilized and stored at -20°C for future use. The molecular weight of the lyophilized α-synuclein was confirmed by mass spectrometry.

To generate the α-synuclein fibril seeds the lyophilised monomeric protein was rehydrated in 1X PBS at a concentration of 500 μM and filtered through a 0.22 μm syringe filter. 500 μl of the filtered protein was transferred to a 2 ml glass vial (27267-U; Sigma-Aldrich) and a sterile magnetic stirrer bar added before the vial was sealed. The vial was then placed inside a mineral oil bath on top of a magnetic stirrer with heating (N2400-3010; STARLAB) and stirred for 3 days at 1500 rpm at 40˚C. The resultant α-synuclein fibril seeds were pelleted down by centrifugation at 16,000Xrcf for 1 hour inside a tabletop centrifuge. The protein concentration of the pellet was checked by BCA assay (J63283; Alfa Aesar) and was diluted down to 200 μM monomeric equivalent concentration with 1X PBS, snap frozen and stored at -80˚C for future use.

#### A90C modification and labelling

To generate the Alexa Fluor 594 labelled A90C α-synuclein fibril seeds, the A90C α-synuclein fibril seeds were generated by the same procedure as the wild-type α-synuclein. After 3 days of stirring at 1500 rpm at 40°C, the resultant A90C α-synuclein fibril seeds were collected by centrifugation at 16,000Xrcf for 1 hour. The protein concentration of the pellet was checked by BCA assay and was diluted down to 100 μM monomeric equivalent concentration with 1X PBS. Dithiothreitol (DTT) was added to the fibril seeds to a final concentration of 5 mM and incubated for 30 minutes on bench. After incubation, the fibril seeds were pelleted down by centrifugation at 16,000 X rcf for 40 minutes, the pellet was kept and washed with 1X PBS by pipette aspiration mixing. The pelleting and washing processes were then repeated for 3 times to remove DTT. The resultant pellet was then labelled with the Alexa Fluor 594 dye by incubating with 10 mM Alexa Flour 594 Maleimide (A10256; Thermo Fisher) at 4°C for 16 hours. Unbound Alexa Fluor 594 Maleimide dye was removed by pelleting and washing processes for 3 times to remove the dye, as performed for the removal of DTT. The protein concentration of the Alexa Fluor 594 labelled A90C fibril seed pellet was checked by BCA assay and diluted down to 200 μM monomeric equivalent concentration with 1X PBS, snap frozen and stored at -80˚C for future use.

#### Generation of a macromolecular crowded bath

A 30% (w/v) Bovine Serum Albumin (BSA) PBS was used to mimic the macromolecular crowded environment of the cell interior. To generate 10 ml of the 30% (w/v) BSA PBS, 3 g of BSA (BS9808K; BioServ UK Ltd.), 1 ml of 10X PBS (AM9625; Thermo Fisher) and 6 ml of dd$H_2O$ were mixed and placed on a roller overnight. The electrolyte solution was stored at -20°C and thawed on the day of usage.

#### Atomic Force Microscopy (AFM) of α-synuclein fibril seeds

2 µl of the A90C modified α-synuclein fibril seeds at 100 µM monomeric equivalent concentration was mixed with 200 µl of 1 M $MgCl_2$ and incubated for 10 minutes on the bench. The mixture was then deposited directly onto a freshly cleaved mica discs and left to incubate for 30 minutes, the samples were then rinsed with PBS for 3 times, 200 µl of PBS was added to the mica disc as imaging buffer. The amyloid fibril samples were imaged using a Bruker Dimension Fastscan (Santa Barbara, CA, USA) with ScanAsyst-Fluid+ cantilevers. The imaging was carried out via PeakForce tapping with ScanAsyst™ liquid imaging mode via Nanoscope software. All images acquired have a pixel resolution of 1024 X 1024, images were analysed with Nanoscope analysis 1.9. The height channel was recorded and analysed with FiberApp [1].

### Supporting figures

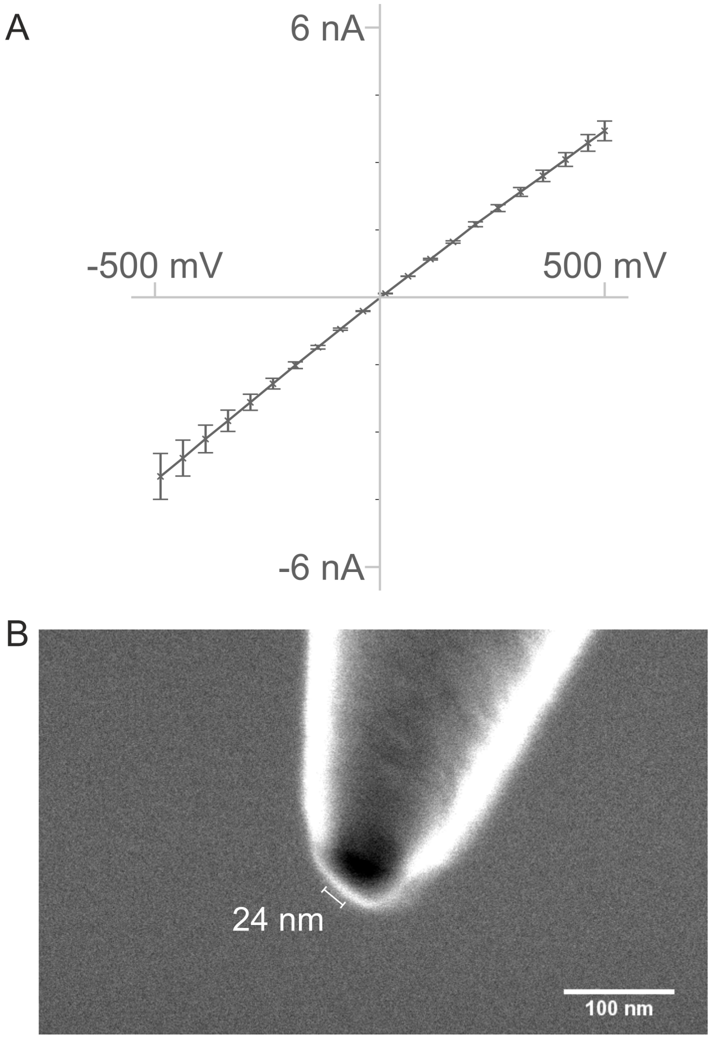


**Supporting Figure 1. Characterisation of the nanopipette.** (A) The average current-voltage (I-V) curve of three nanopipettes (n = 3). Error bars are standard error of the means. (B) The scanning electron micrograph of a nanopipette from (A), it had a pore size of 24 nm.

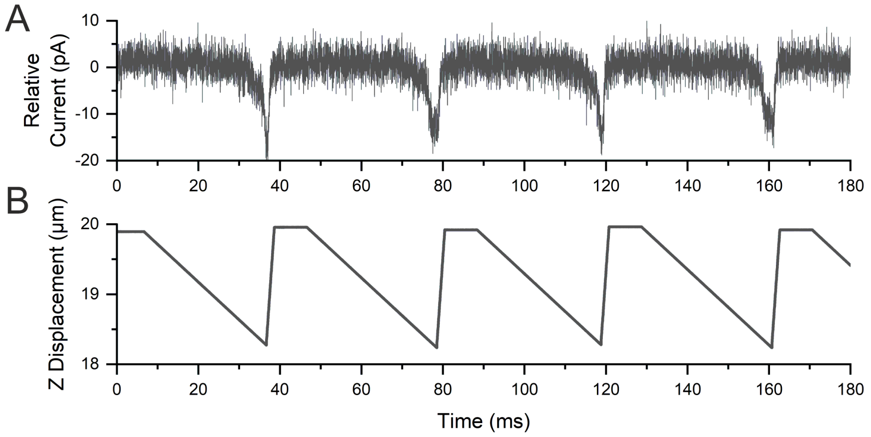


**Supporting Figure 2. The typical hopping trace of the SICM on top of the glass substrate.** The sudden drop in the current (A) occurred when the Z-piezo moved further downward (B). The current returned to baseline when the nanopipette was retracted back to the starting position. Each hop takes approximately 40 ms in this case.

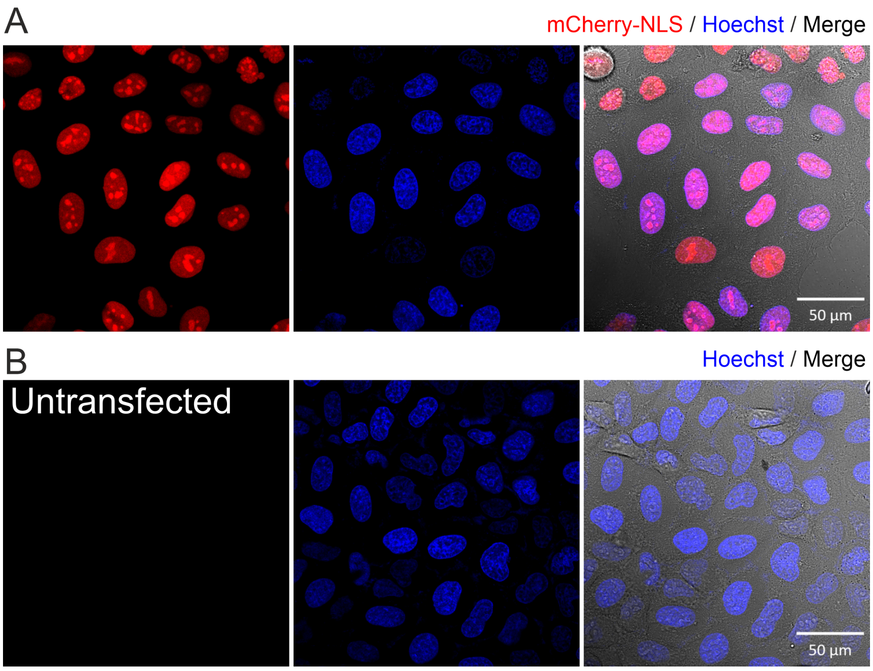


**Supporting Figure 3. The generation of the mCherry-NLS expressing HeLa cell line HeLa RNuc.** The pmCherry-NLS plasmid was transfected to the HeLa cells via lipofection and the HeLa cells were FACS sorted and imaged with a confocal microscope. The pmCherry-NLS transfected cells showed red nuclei co-localized with the Hoechst nuclei staining (A). The untransfected control showed no mCherry-NLS signal (B). Red, mCherry-NLS; Blue, Hoechst 33342; Grey, brightfield image.

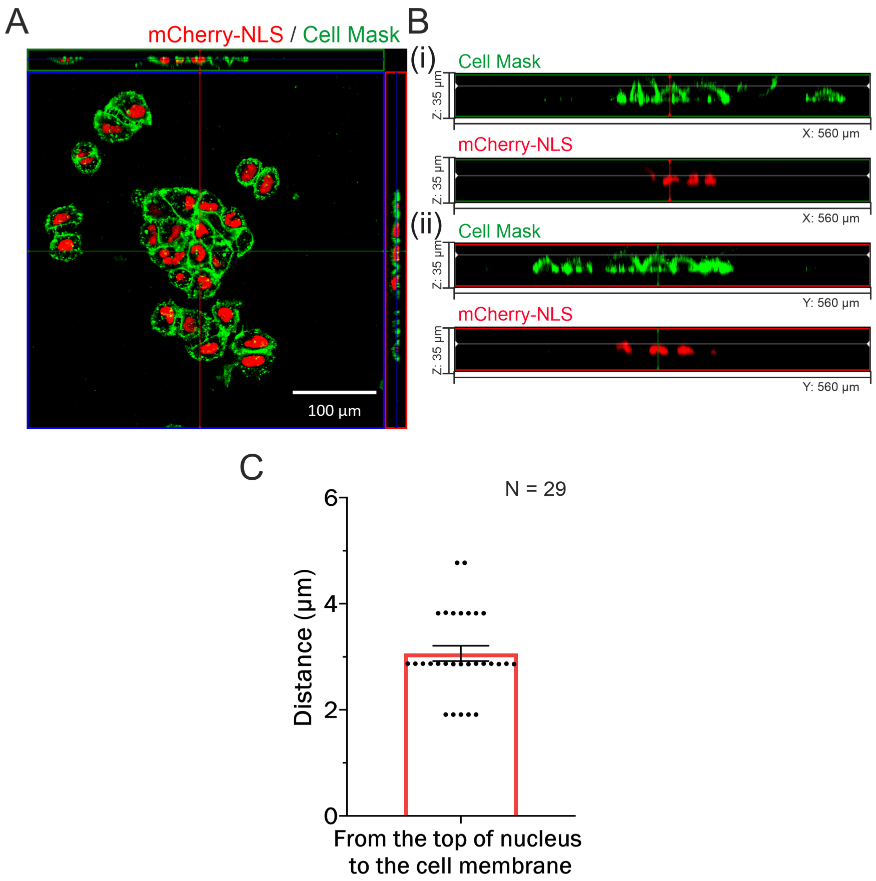


**Supporting Figure 4. The depth of HeLa RNuc nuclei was determined by fluorescent Z-stack imaging.** The orthogonal projection image of the HeLa RNuc cells stained with the Cell Mask$^{TM}$ plasma membrane stains and Hoechst nuclei stain (A). Fluorescent Z-stack imaging was used, 25 stacks were generated for each image at 955.4 nm thickness. The X-Z (i) and X-Y (ii) planes were used to calculate the distance between the top of the cell membrane (Green) and the top of the nucleus (Red) (B). The distance between the top of the cell membrane and the top of the nucleus had an average of 3.1±0.1 µm (N=29 cells). Red, mCherry-NLS; Green, Cell MaskTM green plasma stain.

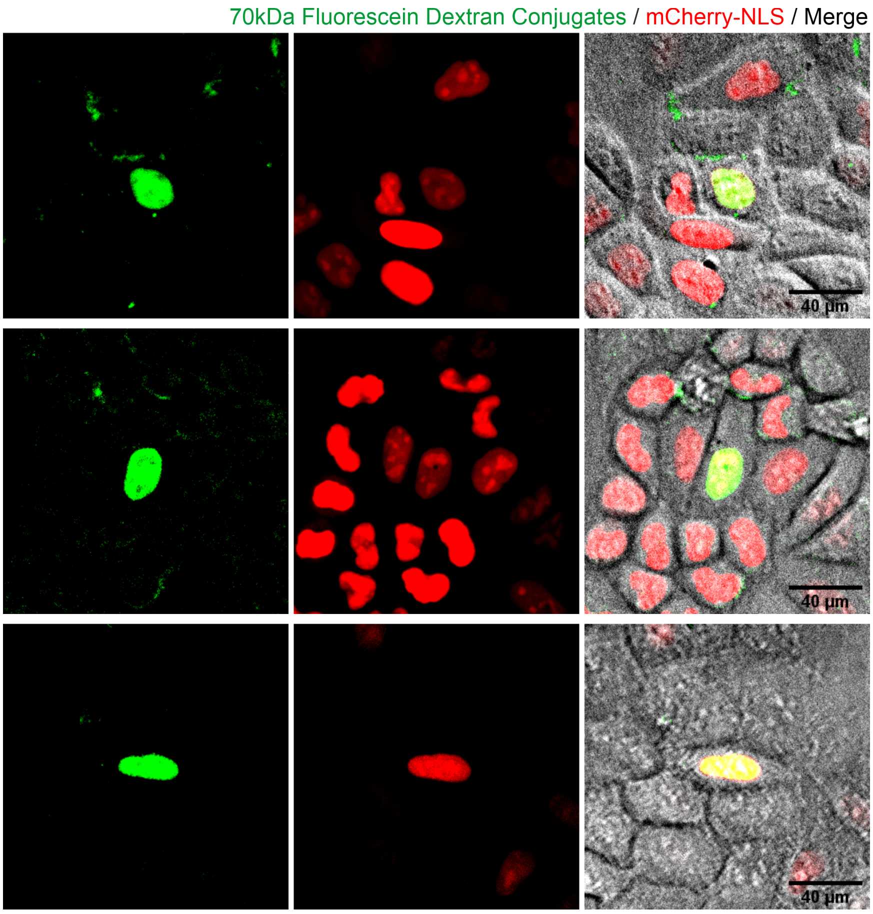


**Supporting Figure 5. The delivery of 70 kDa Fluorescein dextran conjugates into the nuclei of the HeLa RNuc cells.** The 70 kDa fluorescein dextran conjugates were delivered into the nuclei of the HeLa RNuc cells. The size of the conjugates prevented their diffusion into the cytoplasm and were retained inside the nuclei after nuclei nanoinjection. Red, mCherry-NLS; Green, 70 kDa Fluorescein dextran conjugates; Grey, Brightfield image.

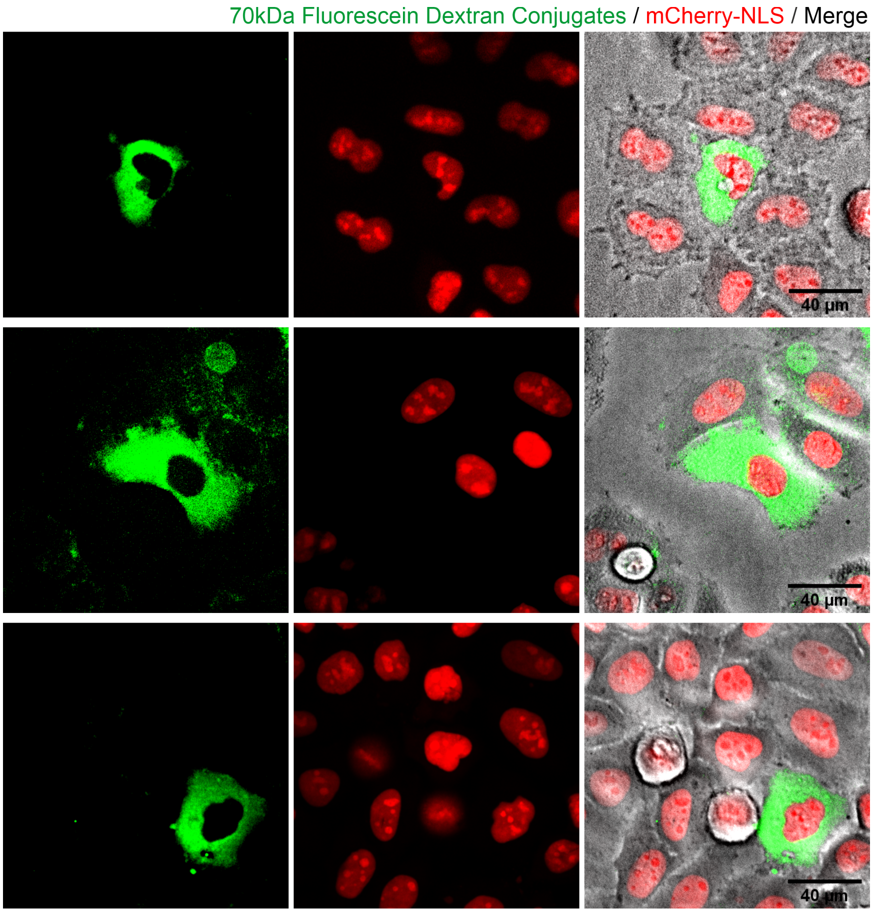


**Supporting Figure 6. The delivery of 70 kDa Fluorescein dextran conjugates into the cytoplasm of the HeLa RNuc cells.** The 70 kDa fluorescein dextran conjugates were delivered into the cytoplasm of the HeLa RNuc cells, the size of the conjugates prevented them from diffusing into the nuclei. Red, mCherry-NLS; Green, 70 kDa Fluorescein dextran conjugates; Grey, Brightfield image.

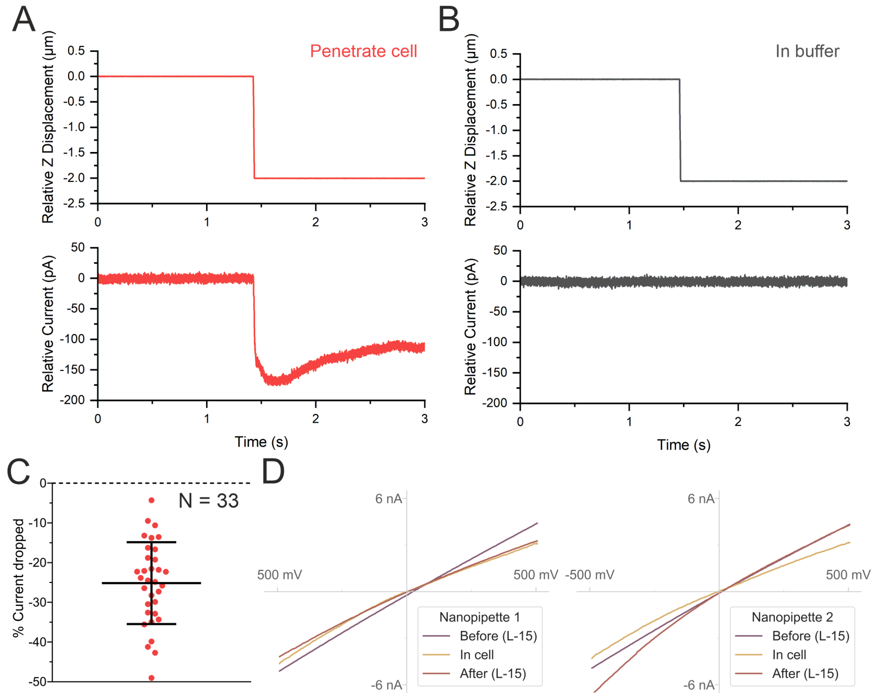


**Supporting Figure 7. Characterising the changes in the ionic current during the penetration of the cell membrane.** The Z-piezo actuator was moved down by 2 μm from the cell membrane to penetrate into the cell (A) or moved down the same distance in pure electrolyte (B). The ionic current initially dropped to about 150 pA when the nanopipette was lowered to penetrate the cell membrane, then the ionic current raised by c.a. 50 pA and stabilised as the new baseline. The percentage of the drop in ionic current between the pre-penetration baseline and the post-penetration baseline showed an average of 25.2 ± 1.8% drop (N=33) (C). The IV curves of two nanopipettes in the L-15 medium, inside cell, and after nanoinjection in L-15 medium (D), the slight deviation in the IV profile may be due to the intracellular environment and cellular materials sticking to the nanopipette wall. The similar levels of currents suggested that the nanopore is not damaged by the penetration of the cell.

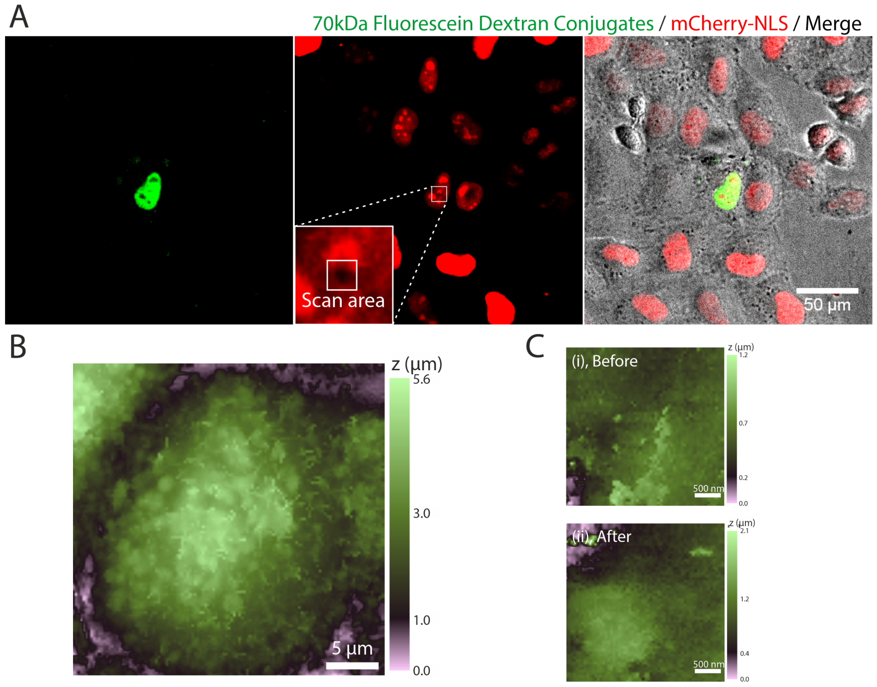


**Supporting Figure 8. Spatial control of the nanoinjection of HeLa cells enabled by SICM.** (A) The nuclear nanoinjection of 70 kDa fluorescein dextran conjugate to the mCherry-NLS expressing HeLa cells, the dye fluorescent signal was localized and overlapped with the mCherry-NLS signal. (B) Topography maps acquired using the SICM of the HeLa cells from (A). The zoom in scan of the highlighted black dot area from (A), revealed no substantive membrane deformation and no hole in the plasma membrane. (i) Before the nanoinjection and (ii) after the nanoinjection.

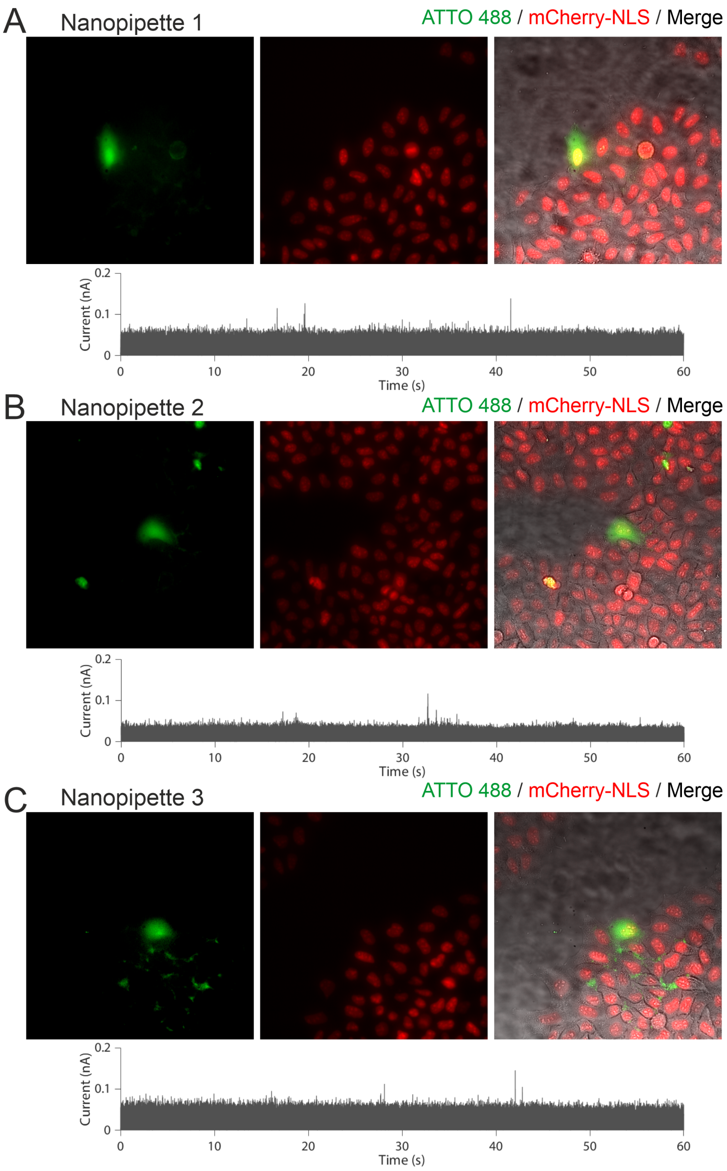


**Supporting Figure 9. Current traces inside HeLa-RNuc cells for 10 µM ATTO 488 1X PBS filled nanopipettes.** The 10 µM ATTO 488 1X PBS injection trace inside three cells (A, B & C). Less than 10 peaks were observed may be due to the translocation of intracellular materials into the nanopipettes.

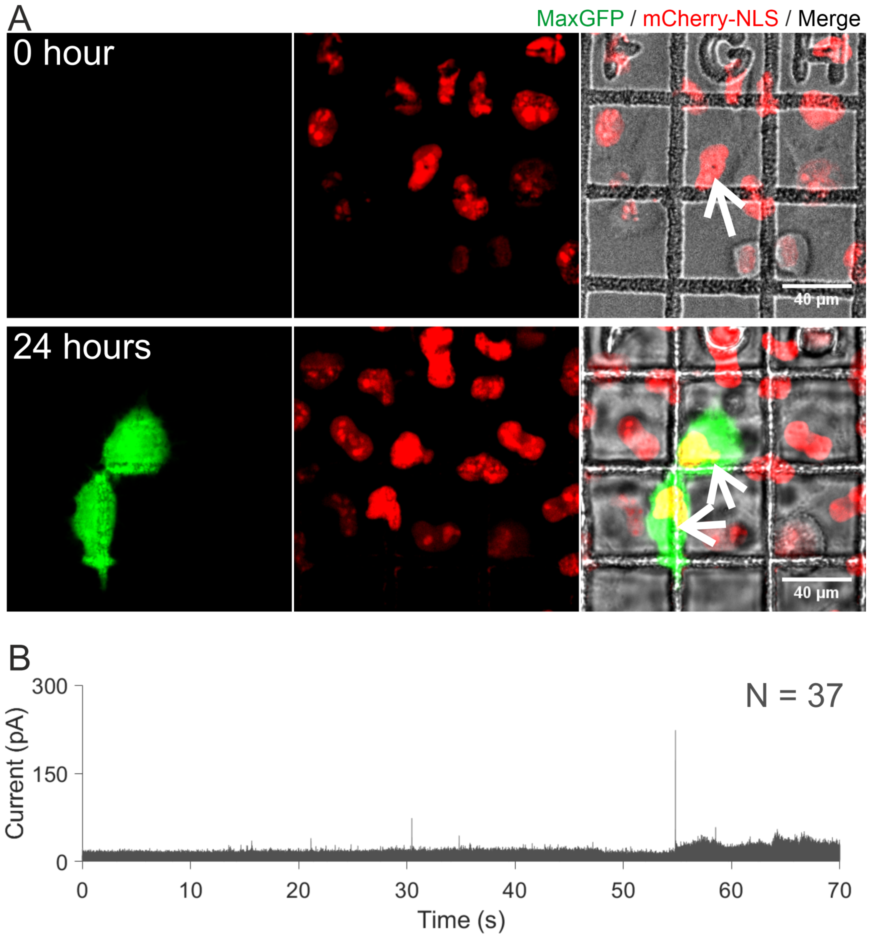


**Supporting Figure 10. Single cell transfection of the HeLa RNuc cells with the pMaxGFP via quantitative nanoinjection – 2.** HeLa RNuc cells were plated onto a gridded glass dish (50 X 50 µm per square). On the day of the nanoinjection (0 hours), a cell was nanoinjected with the pMaxGFP (arrow) and the same cell was imaged 24 hours later with the aid of the grid, the cell expressed MaxGFP and had divided (arrows) (A). The baseline adjusted ionic current trace during the nanoinjection was recorded and analysed, and a total of 37 translocation events were detected during nanoinjection (B).

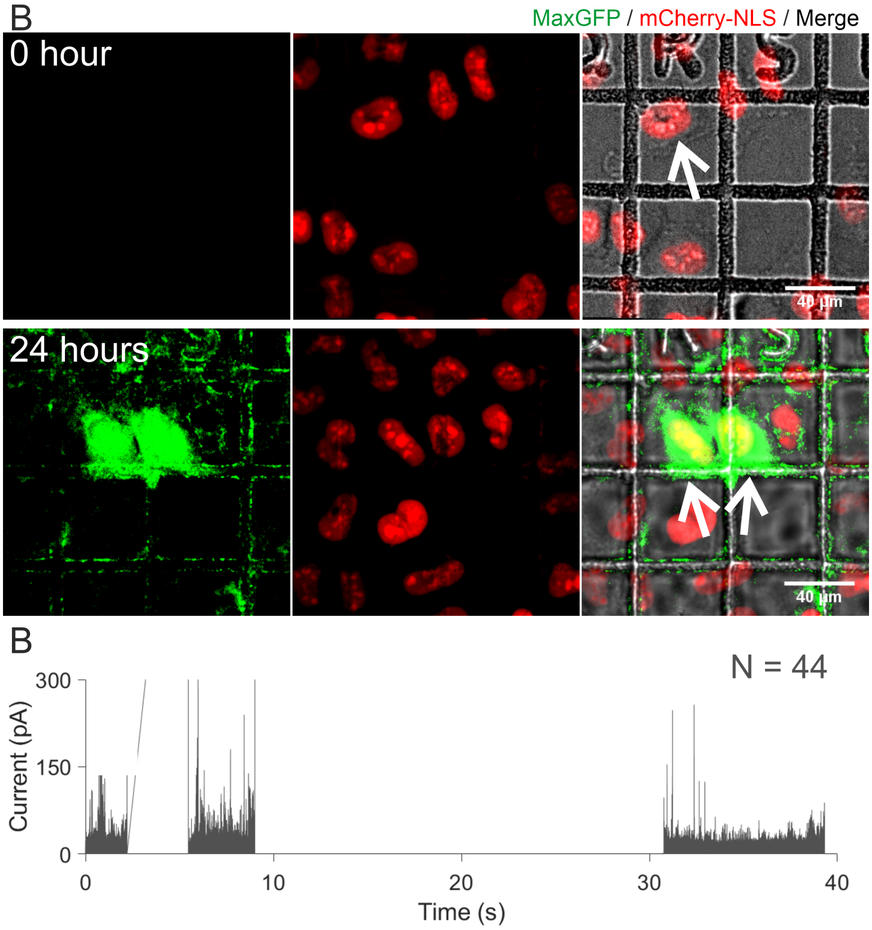


**Supporting Figure 11. Single cell transfection of the HeLa RNuc cells with the pMaxGFP via quantitative nanoinjection – 3.** HeLa RNuc cells were plated onto a gridded glass dish (50 X 50 µm per square). On the day of the nanoinjection (0 hours), a cell was nanoinjected with the pMaxGFP (arrow) and the same cell was imaged 24 hours later with the aid of the grid, the cell expressed MaxGFP and had divided (arrows) (A). The baseline adjusted ionic current trace during the nanoinjection was recorded and analysed, and a total of 44 translocation events were detected during nanoinjection (B). Gaps in the ion current traces are caused by fluctuations in the signal preventing robust data analysis.

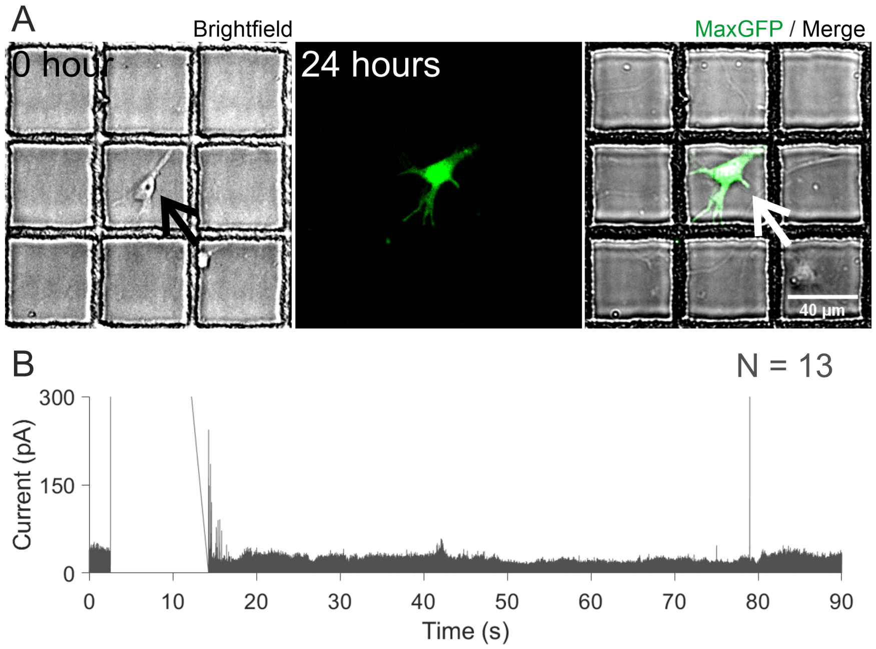


**Supporting Figure 12. Single cell transfection of a primary DRG neuron with the pMaxGFP via quantitative nanoinjection – 2.** Primary DRG neurons were plated onto a gridded glass dish (50 X 50 µm per square). On the day of the nanoinjection (0 hours), a cell was nanoinjected with the pMaxGFP (arrow), and the same cell was imaged 24 hours later with the aid of the grid. The cell expressed MaxGFP (A). The baseline adjusted ionic current trace during the nanoinjection was recorded and analysed, and a total of 13 translocation events were detected during nanoinjection (B). Gaps in the ion current traces are caused by fluctuations in the signal preventing robust data analysis.

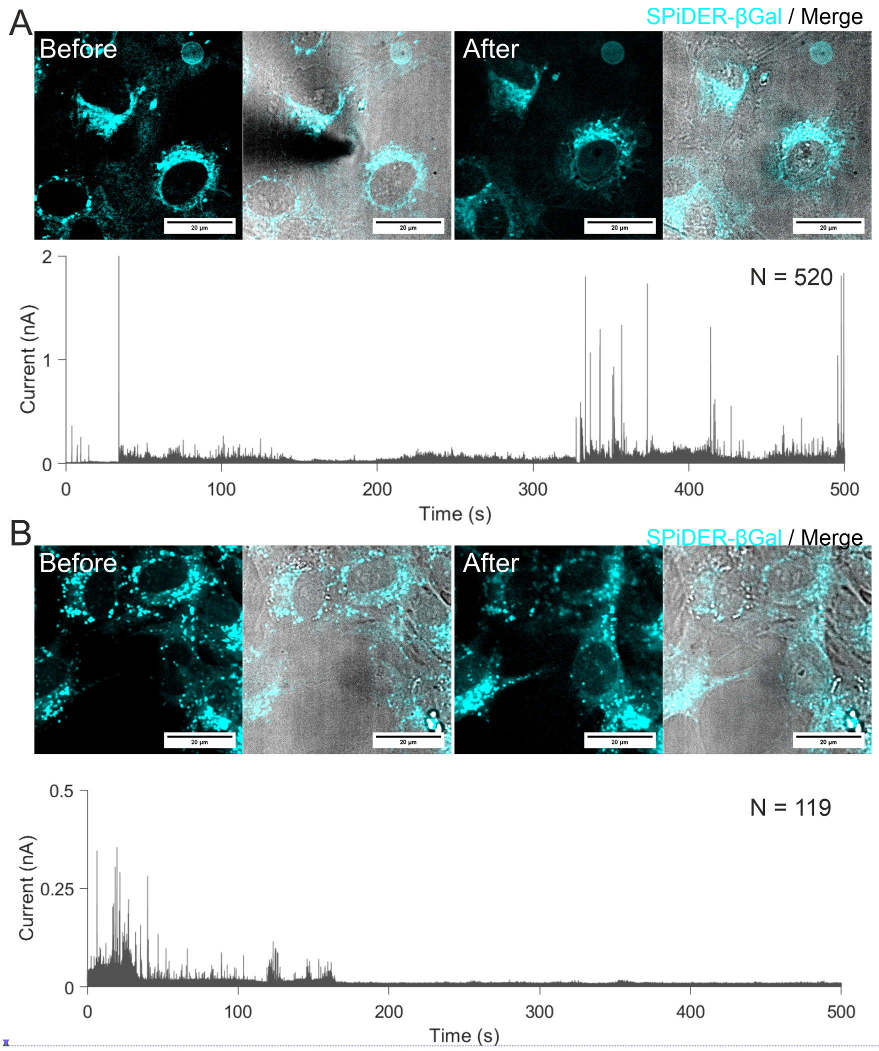


**Supporting Figure 13. Intracellular quantitative nanoinjection of the enzyme β-galactosidase – 1&2.** Purified β-galactosidase diluted to 1 μM in PBS was used to fill the nanopipette. The SPiDER-βGal was added to the SICM imaging buffer at a final concentration of 2 μM. Fluorescent and brightfield images of the SPiDER-βGal channel and brightfield were taken before the nanoinjection. Once the nanopipette was inside the cell, a voltage of -700 mV was used to deliver the β-galactosidase into the nucleus. The fluorescent and brightfield images were taken after the nanoinjection. 520 molecules (A) and 110 molecules (B) were recorded from the current trace during the injection.

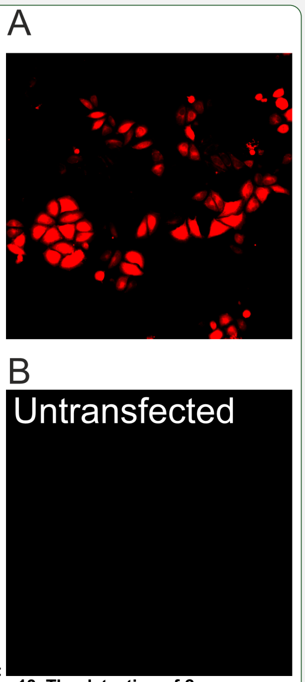


**Moved down [1]:** **Supporting Figure 13. The detection of β-galactosidase activity in cells using the fluorescent substrate SPiDER-βGal.** The HeLa cells were transfected with the commercially available pSV-β-galactosidase plasmid with the lipofectamine 2000. After the transfection, the cells were incubated with 2 μM of SPiDER-βGal for 15 minutes and then imaged by a widefield fluorescence microscope. The pSV-β-galactosidase transfected cells exhibited red fluorescence after the incubation with the SPiDER-βGal and excitation with suitable wavelength light (A). The untransfected control showed no red fluorescent signal (B). Red, SPiDER-βGal; Blue, Hoechst 33342; Grey, brightfield image.

Page Break

**Deleted:**

Page Break

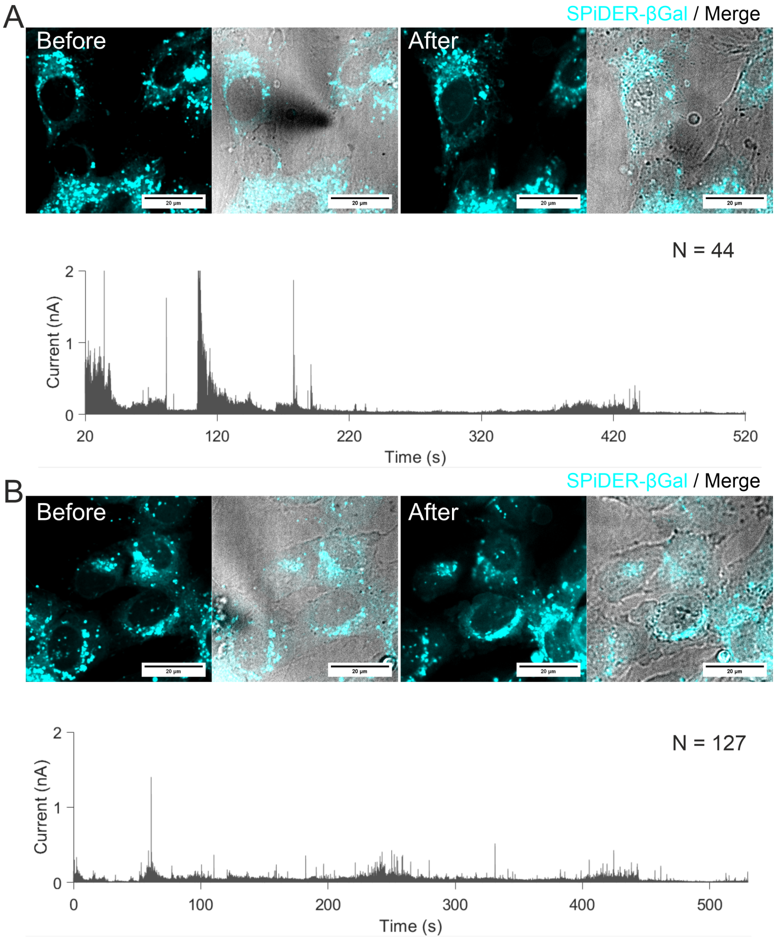


**Supporting Figure 14. Intracellular quantitative nanoinjection of the enzyme β-galactosidase – 3&4.** Purified β-galactosidase - diluted to 1 μM in PBS was used to fill the nanopipette. The SPiDER-βGal was added to the SICM imaging buffer at a final concentration of 2 μM. Fluorescent and brightfield images of the SPiDER-βGal channel and brightfield were taken before the nanoinjection. Once the nanopipette was inside the cell, a voltage of -700 mV was used to deliver the β-galactosidase into the nucleus. The fluorescent and brightfield images were taken after the nanoinjection. 44 molecules (A) 127 molecules (B) were recorded from the current trace during the injection.

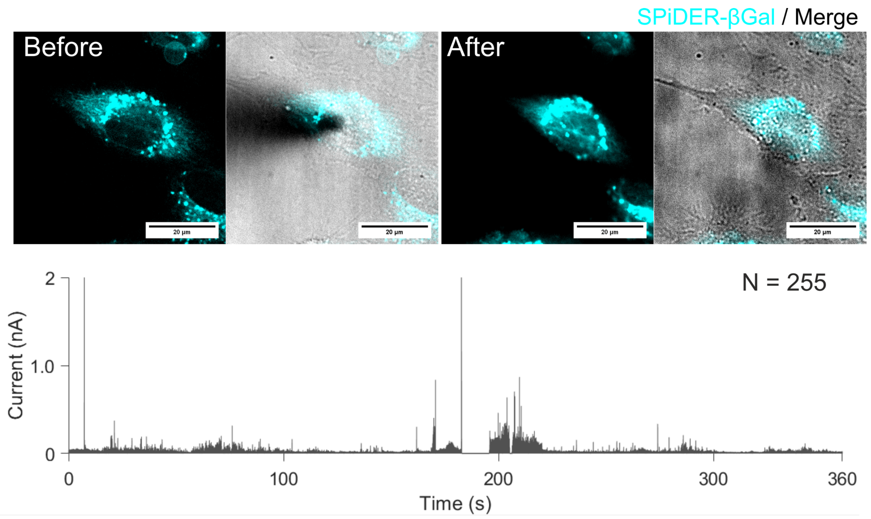


**Supporting Figure 15. Intracellular quantitative nanoinjection of the enzyme β-galactosidase – 5.** Purified β-galactosidase diluted to 1 μM in PBS was used to fill the nanopipette. The SPiDER-βGal was added to the SICM imaging buffer at a final concentration of 2 μM. Fluorescent and brightfield images of the SPiDER-βGal channel and brightfield were taken before the nanoinjection. Once the nanopipette was inside the cell, a voltage of -700 mV was used to deliver the β-galactosidase into the nucleus. The fluorescent and brightfield images were taken after the nanoinjection. 255 molecules were recorded from the current trace during the injection.

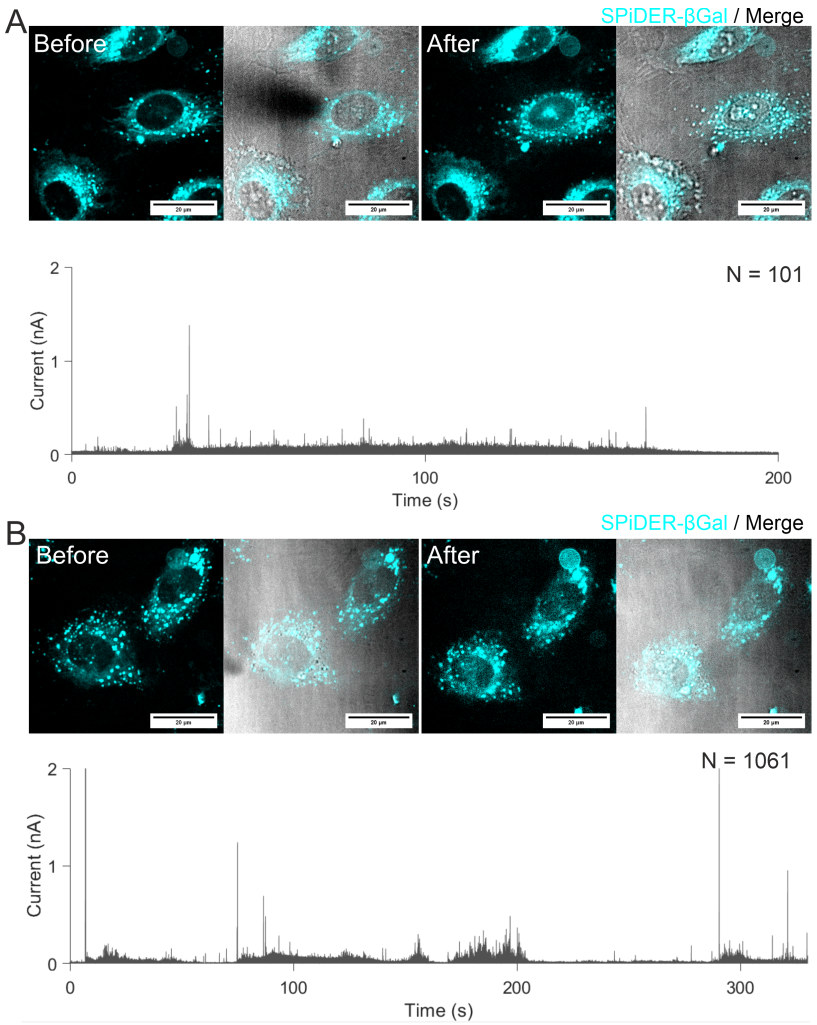


**Supporting Figure 16. Intracellular quantitative nanoinjection of the enzyme β-galactosidase – 6&7.** Pβ-galactosidase =diluted to 1 μM in PBS was used to fill the nanopipette. The SPiDER-βGal was added to the SICM imaging buffer at a final concentration of 2 μM. Fluorescent and brightfield images of the SPiDER-βGal channel and brightfield were taken before the nanoinjection. Once the nanopipette was inside the cell, a voltage of -700 mV was used to deliver the β-galactosidase into the nucleus. The fluorescent and brightfield images were taken after the nanoinjection. 101 molecules (A) 1061 molecules (B) were recorded from the current trace during the injection.

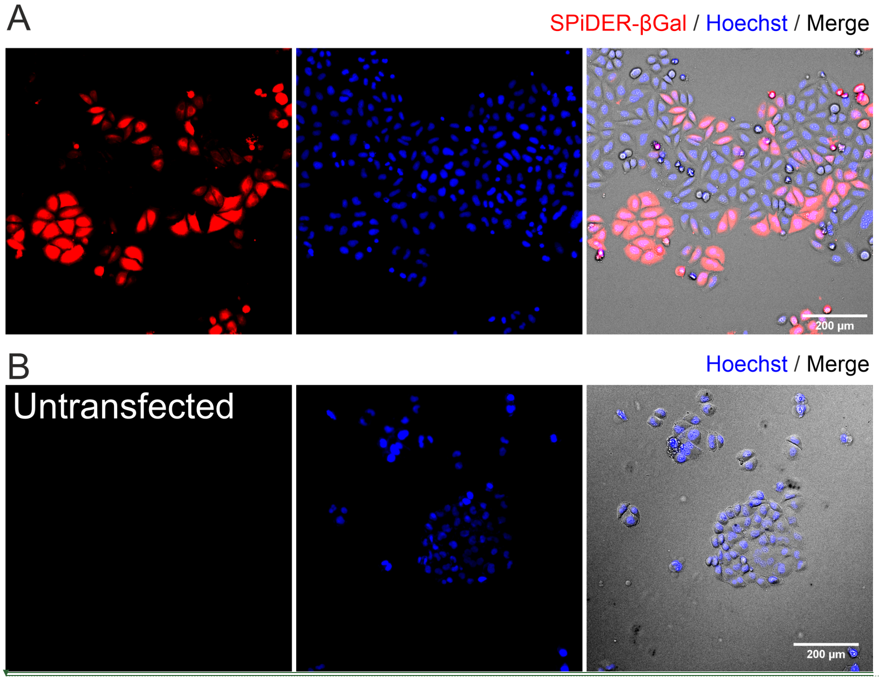


**Supporting Figure 17. The detection of β-galactosidase activity in cells using the fluorescent substrate SPiDER-βGal.** The HeLa cells were transfected with the commercially available pSV-β-galactosidase plasmid with the lipofectamine 2000. After the transfection, the cells were incubated with 2 μM of SPiDER-βGal for 15 minutes and then imaged by a widefield fluorescence microscope. The pSV-β-galactosidase transfected cells exhibited red fluorescence after the incubation with the SPiDER-βGal and excitation with suitable wavelength light (A). The untransfected control showed no red fluorescent signal (B). Red, SPiDER-βGal; Blue, Hoechst 33342; Grey, brightfield image.

**Moved (insertion) [1]**

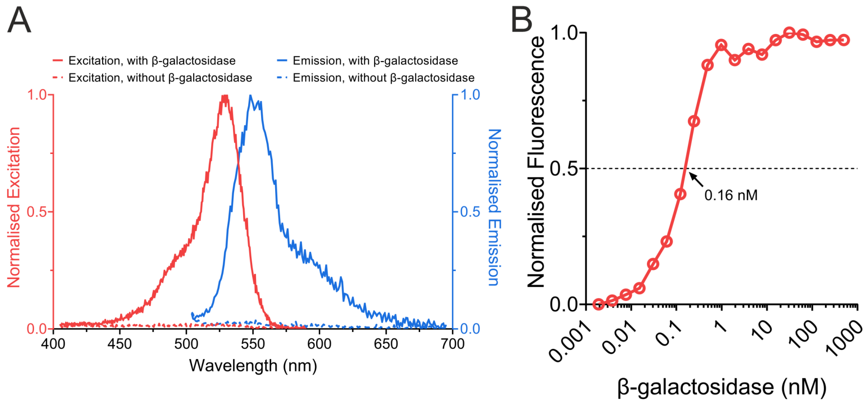


**Supporting Figure 18. The fluorescence excitation/emission spectra of SPiDER-βGal before and after hydrolysis by β-galactosidase and the sensitivity of SPiDER-βGal for the detection of β-galactosidase activity in solution.** 1 μM β-galactosidase was incubated with 2 μM SPiDER-βGal and the emission/excitation spectra analysed. The excitation maximum is at 530 nm and emission maximum is at 550 nm. Without the addition of the SPiDER-βGal substrate, there is a minimal fluorescent signal (A). The sensitivity of the fluorescent substrate in the measurement of β-galactosidase activity was analysed by performing a serial dilution of the β-galactosidase from 1 μM down to 1 pM. 2 μM of SPiDER-βGal substrate was added to the β-galactosidase and incubated for 5 minutes at room temperature prior to measuring the fluorescent signal with a plate reader. 0.16 nM of β-galactosidase was required to obtain half of the maximum fluorescent signal (B).

**Formatted:** Normal, Line spacing: single

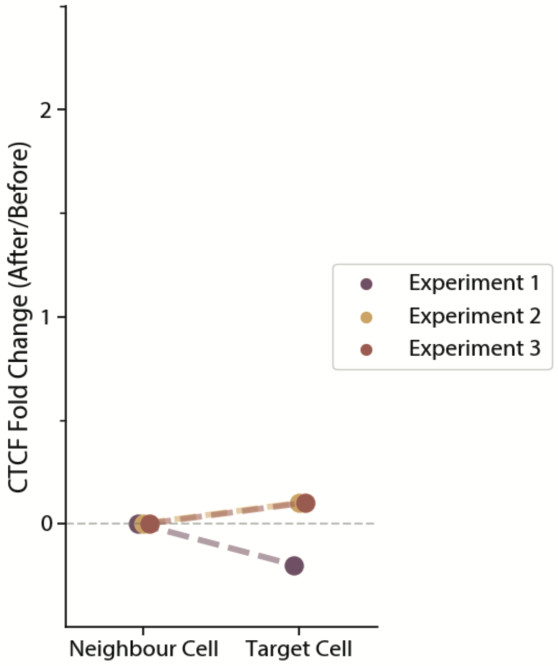


**Supporting Figure 19. The Corrected Total Cell Fluorescence (CTCF) of the nucleus area before and after the nanoinjection.** The cells were nanoinjected with PBS buffer, the CTCF fold changes before/after injection comparison between the target nanoinjected cell and a neighbouring control cell.

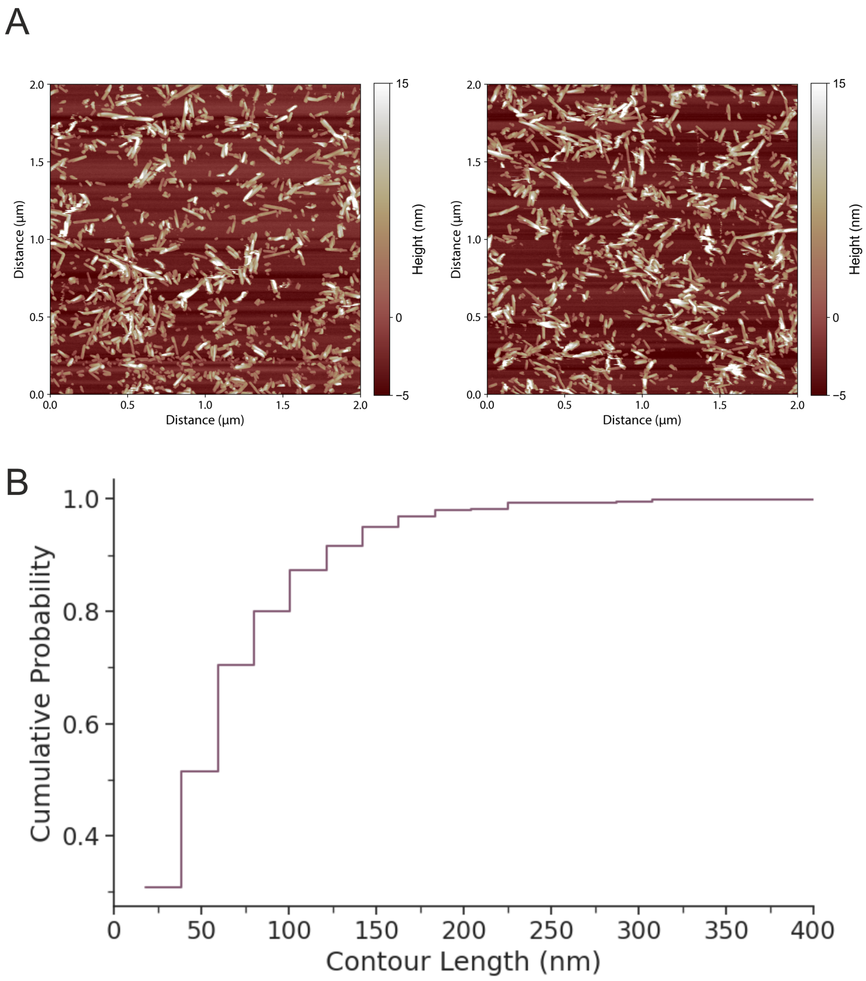


**Supporting Figure 20. AFM characterisation of the Alexa fluor 594 labelled A90C α-synuclein fibrils.** (A) AFM images of the fluorescently labelled A90C α-synuclein fibrils. (B) Cumulative histogram showing that 80% of the fibrils have a length below or equal to 100 nm.

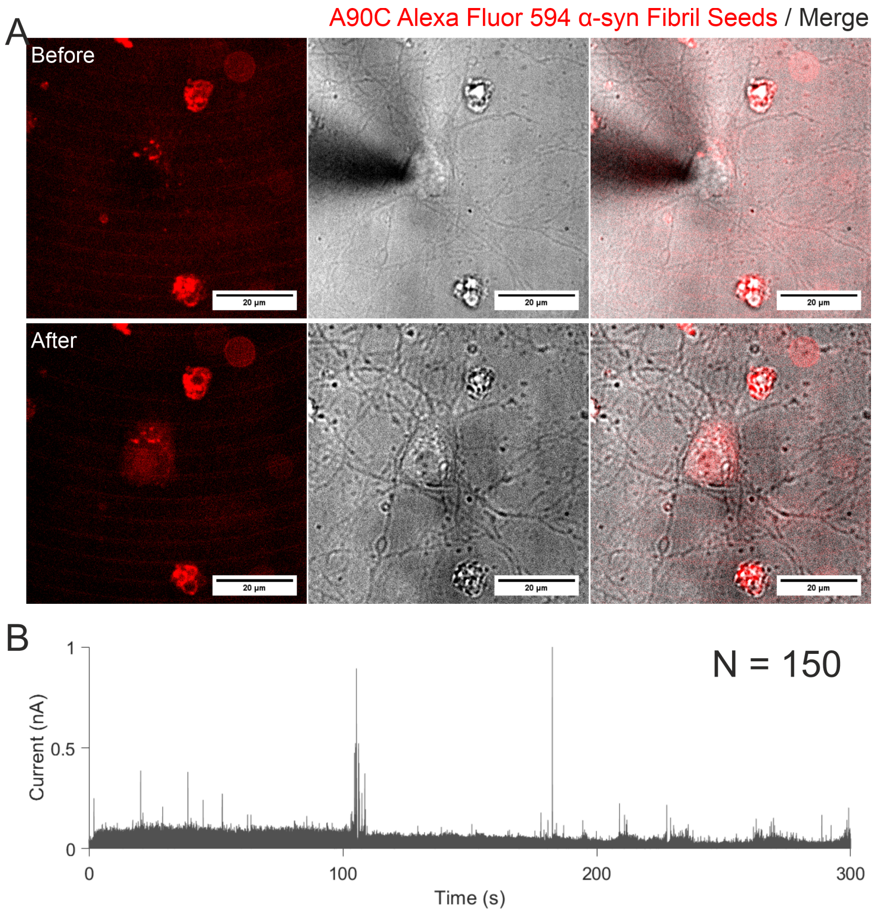


**Supporting Figure 21. Quantitative nanoinjection of fluorescently labelled α-synuclein fibrils into primary rat cortical neurons - 1.** The nanopipette was filled with 1 µM monomeric equivalent A90C Alexa Fluor 594 labelled α-synuclein fibrils diluted in PBS. (A) The images before and after nanoinjection of the neurons, revealed that the neurons exhibited increased fluorescence after nanoinjection with the fluorescently labelled fibrils. (B) Ion current traces corresponding to the injections. The molecule count and the trace showed 150 molecules were detected.

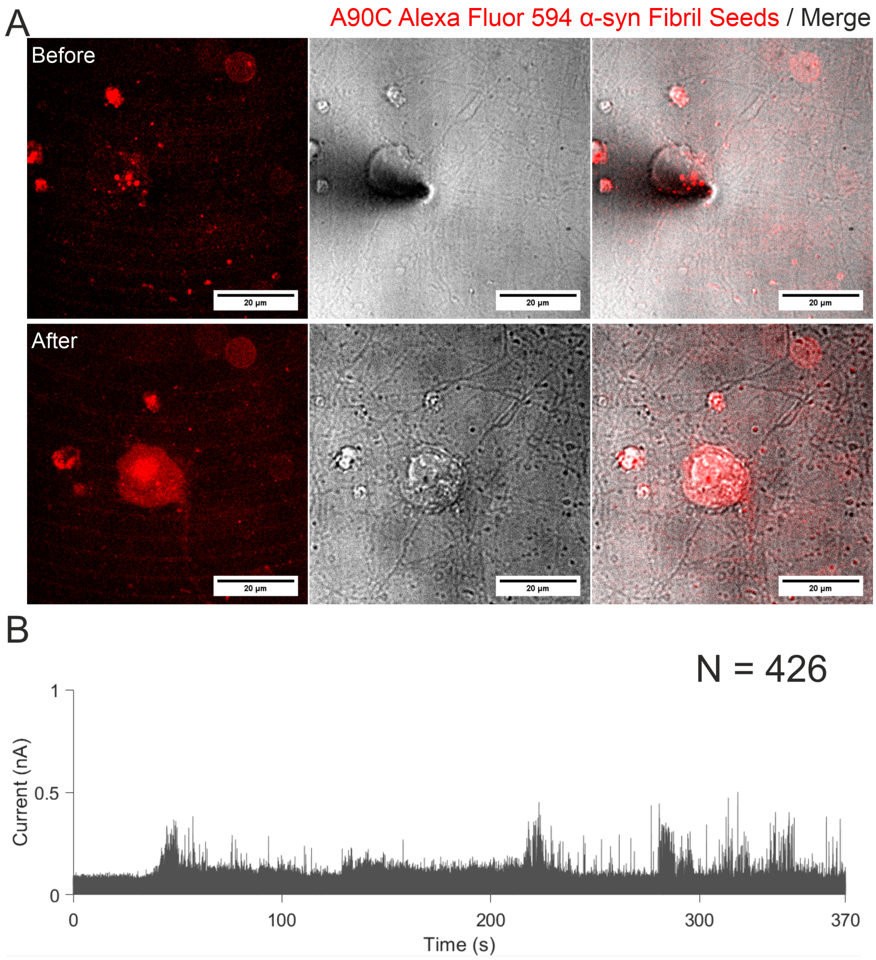


**Supporting Figure 22. Quantitative nanoinjection of fluorescently labelled α-synuclein fibrils into primary rat cortical neurons - 2.** The nanopipette was filled with 1 µM monomeric equivalent A90C Alexa Fluor 594 labelled α-synuclein fibrils diluted in PBS. The images before and after nanoinjection of the neurons, revealed that the neurons exhibited increased fluorescence positive after nanoinjection with the fluorescently labelled fibrils (A). The molecule count and the trace showed 426 molecules were detected (B).

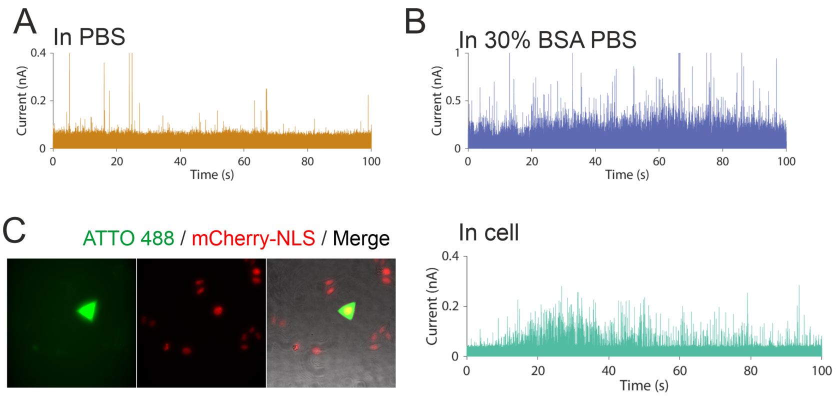


**Supporting Figure 23. The translocation of the 7 kbp dsDNA from the same nanopipette into three different external environments.** A 5 nM 7 kbp dsDNA in 10 μM ATTO 488 PBS was nanoinjected into PBS (A), 30% BSA PBS (B) and into a HeLa RNuc cell (C). The cell became fluorescent after the nanoinjection of the dsDNA dye mixture, confirming that the translocation peaks observed were from nanoinjection into the intracellular environment.

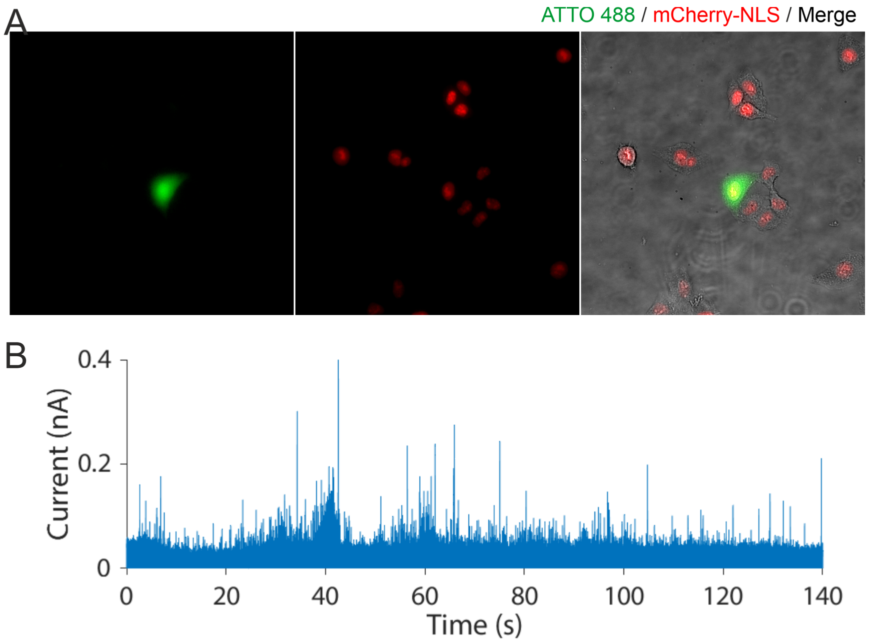


**Supporting Figure 24. The translocation of the 7 kbp dsDNA into a cell - 2.** The 5 nM 7 kbp dsDNA in 10 μM ATTO 488 PBS was nanoinjected into a HeLa-RNuc cell. The fluorescent images confirmed that the translocation of the nanopipette's contents into the cell was carried out (A), and the associated translocation trace of the dsDNA (B).

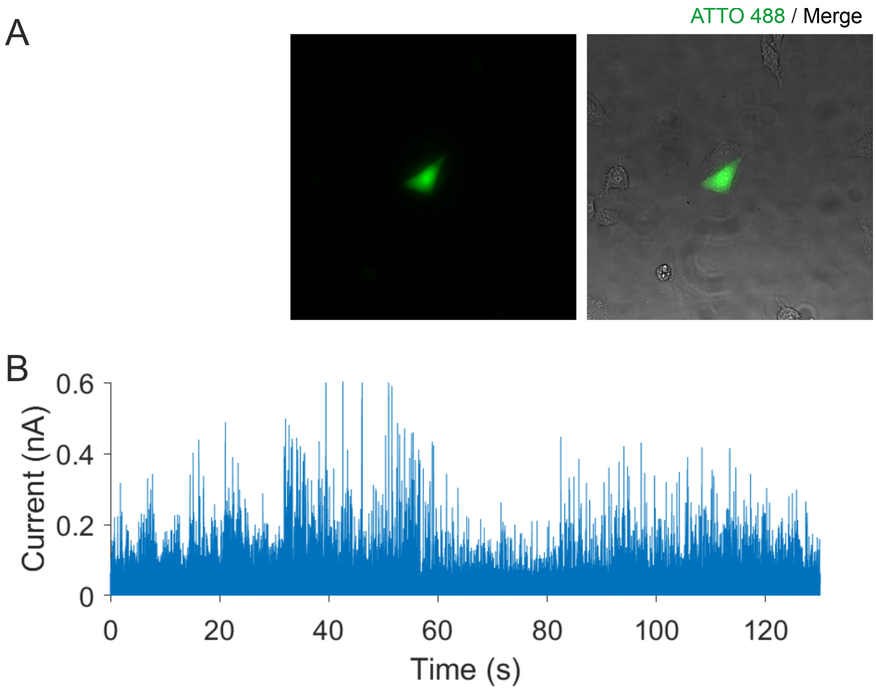


**Supporting Figure 25. The translocation of the 7 kbp dsDNA into the a - 3.** The 5 nM 7 kbp dsDNA in 10 µM ATTO 488 PBS was translocated into a HeLa cell. The fluorescence images confirmed that the translocation of the nanopipette contents into the cell was carried out (A), and the associated translocation trace of the dsDNA (B).

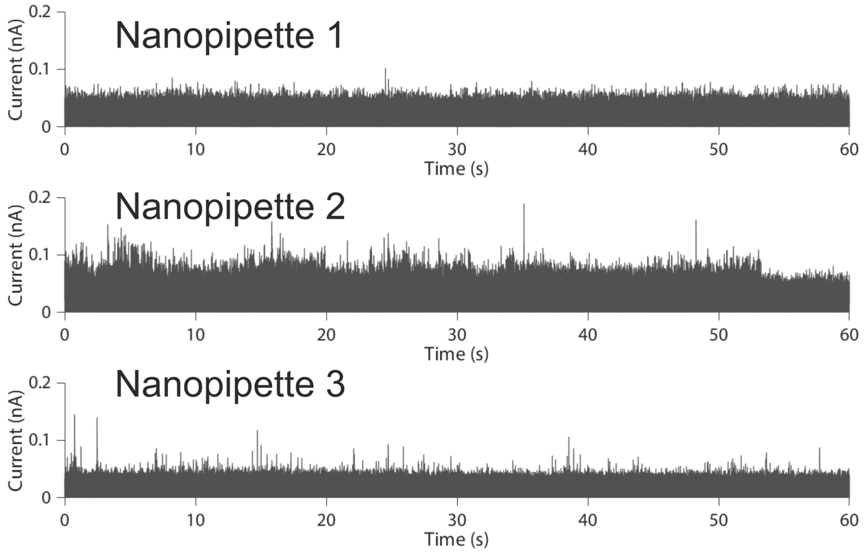


**Supporting Figure 26. The 1X PBS filled nanopipette inside the 30% BSA PBS bath.** The current traces of three nanopipettes were analysed at -500 mV in a 30% BSA PBS bath. The peaks observed could be due to the translocation of BSA into the nanopipette.
